# Differential responses of murine embryonic stem cells (mESC) and their endothelial progeny to doxorubicin and pharmacological inhibitors of DNA repair and DNA damage response

**DOI:** 10.1101/2025.05.15.654253

**Authors:** Sina Federmann, Michelle Westerhoff, Andreas S. Reichert, Gerhard Fritz

**Affiliations:** Institute of Toxicology, Medical Faculty and University Hospital, Heinrich-Heine-University Duesseldorf, Moorenstrasse 5, 40225 Duesseldorf, Germany; Institute of Biochemistry and Molecular Biology I, Medical Faculty and University Hospital, Heinrich-Heine-University Duesseldorf, Universitätsstrasse 1, 40225 Duesseldorf, Germany

**Keywords:** Murine embryonic stem cells (mESC), endothelial differentiation, anthracyclines, DNA damage, DNA repair, DNA damage response (DDR), cardiotoxicity, regeneration, developmental toxicology

## Abstract

The clinical use of the anticancer drug doxorubicin (Dox) is limited by irreversible cardiotoxicity. The detailed molecular mechanisms involved and the pathophysiological relevance of different cardiac cell types, including progenitor cells, are still unclear. Here, we investigated stress responses of murine embryonic stem cells (mESC), endothelial progenitor cells (EC d4) and terminally differentiated endothelial-like cells (EC d6) following exposure to Dox and selected pharmacological inhibitors of DNA repair and DNA damage response (DDR) (RAD51i B02 and HDACi entinostat (EST)). We found that EC d4 exhibited a pronounced Dox hypersensitivity as compared to both mESC and EC d6, which was independent of drug transport. Analysis of EdU incorporation and replication fork progression following drug treatment revealed substantial agent-specific differences between mESC and the various differentiation stages. Furthermore, cellular susceptibility to drug-induced formation of DNA damage (i.e. DSB and SSB) also changes with ongoing differentiation and drug treatment, with mESC and EC d4 / EC d6 being particular prone to enhanced residual SSB and DSB levels, respectively. Dox treatment of EC d4 did not affect their differentiation into EC d6, but caused multiple functional impairments of the surviving EC d6 progeny, including defects in mitochondrial homeostasis, barrier function related to cell-cell adhesion factors ZO1 and VE-cadherin, response to cytokine stimulation as well as LDL uptake. To summarize, we show substantial differences in the response of mESC, EC d4 and EC d6 to Dox and pharmacological inhibitors of DNA repair and DDR. Most important, treatment of EC d4 results in pronounced persisting functional impairments of differentiated EC d6, pointing to a transient particularly drug-sensitive time window during endothelial differentiation. These findings are important for hazard assessment in developmental toxicology and regenerative medicine in the context of anticancer drug-induced normal tissue damage.

**Highlights:**

- EC d4 are most vulnerable towards Dox-induced cytotoxicity independent of drug transport
- EC d4 and EC d6 display higher steady-state levels of drug-induced DSB as compared to mESC
- Drug-induced replication stress is highest in mESC and decreases with differentiation.
- Both Dox and DNA repair/DDR inhibitors damage mitochondrial homeostasis in EC d4 and EC d6
- Terminally differentiated EC d6 derived from drug-treated EC d4 display multiple functional impairments

## 1. Introduction

Anthracycline derivatives such as doxorubicin (Dox) are widely used anticancer drugs [1], which act as type II topoisomerase (Topo II) poisons, thereby leading to the formation of highly cytotoxic DNA double-strand breaks (DSBs) [2]. DSBs are highly potent triggers of the DNA damage response (DDR), which regulates cell cycle progression, DNA repair and apoptosis, thereby defining the balance between survival and death-related mechanisms [3]. In addition, the formation of reactive oxygen species (ROS), DNA intercalation, inhibition of helicases and chromatin damage contribute to Dox-induced cytotoxicity [4, 5]. Irreversible cardiotoxicity ultimately leading to cardiomyopathy and congestive heart failure is the clinically most relevant adverse effect of anthracyclines [1]. In view of the low antioxidative capacity of cardiomyocytes, mitochondria-related iron-dependent and -independent ROS formation might be of pathophysiological relevance for Dox-induced cardiac damage, with p53-regulated mechanisms of senescence and cell death being involved [6–10]. Hence, prevention of anthracycline-mediated oxidative stress has been considered to achieve cardioprotection [9–11]. Yet, antioxidants failed to demonstrate substantial cardioprotective potency following anthracycline-based therapy [4, 12, 13]. By contrast, the EDTA analogue dexrazoxane, which is a strong catalytic inhibitor of topoisomerase II, has been proven to prevent anthracycline-induced cardiac damage [14, 15]. Based on data obtained from the use of dexrazoxane derivatives lacking ion-chelating activity but still harboring Topo II inhibitory potency, it is hypothesized that their cardioprotective activity mainly rests on the inhibition of Topo II, in particular Topo IIβ isoform [16, 17]. This hypothesis is supported by the observation that knockout of Top2β protects from anthracycline-induced cardiotoxicity *in vivo* [16].

Regarding the pathophysiology of anthracycline-induced heart damage it is still unclear which cardiac cell types are of utmost relevance for acute and chronic cardiotoxicity. Both cardiomyocytes, endothelial cells, fibroblasts, immune cells and cardiac progenitor cells have been suggested to contribute to Dox-mediated heart injury [18, 19]. In view of the delayed onset of cardiotoxicity under clinically relevant setting of repeated Dox treatment cycles, it is tempting to speculate that Dox-induced persisting damage to progenitor cells might lead to a further impairment of the already limited regenerative capacity of the heart. In addition, it was recently suggested that the pathophysiological relevance of cardiomyocytes versus non-cardiomyocytes for cardiac damage following anthracycline exposure also depends on the time point of analysis (acute vs. chronic), concrete treatment regimen (single vs repeated treatments) and the dose (single, cumulative) applied [20, 21]. Having in mind the pathophysiological relevance of endothelial damage [18, 22–25] as well as the involvement of cardiac progenitor cells in Dox-induced cardiotoxicity [26–28], our study aims to comparatively investigate the vulnerability of murine embryonic stem cells (mESC), thereof derived endothelial progenitor cells (EC d4) undergoing differentiation and *in vitro* differentiated endothelial-like cells (EC d6) to Dox-induced damage. In view of the putative pathophysiological relevance of DNA double-strand breaks (DSBs), which are resulting from Topo II inhibition, in Dox-triggered cardiotoxicity [17, 29–32], we included pharmacological inhibitors of DSB repair by homologous recombination (HR) and DNA damage response (DDR) into our study. Thereby, we aimed to examine the outcome of DNA damage caused by differentiation-associated endogenous processes. In more detail, cell viability, proliferation, activation of cell death and senescence-associated mechanisms, mitochondrial homeostasis, the steady state levels of DSB and SSB as well as the activation of DDR related mechanism were comparatively analyzed. Most important, the outcome of drug treatment of EC d4 on prototypical endothelial functions of terminally differentiated EC d6 was investigated in order to assess the long-term toxic effects of drug exposure of precursor cells on the fitness of their progeny.

## 2. Materials and Methods

### 2.1 Materials

Mouse ESC (LF2) were isolated from the mouse strain 129J [33] and originate from A. Smith (Oxford, UK). Activin A, BMP4, bFGF, and VEGF165 are from PeproTech (Hamburg, Germany), GSKi from Calbiochem (Darmstadt, Germany), ALKi and entinostat (EST) from Sellek Chemicals LLC (Munich, Germany), Forskolin and Hydroxyurea (HU) from Sigma-Aldrich (Munich, Germany), doxorubicin (Dox) from Cellpharm (Bad Vilbel, Germany) and B02 was from Tocris Bioscience (Bristol, UK). Antibodies originate from the following companies: VE-cadherin from eBioscience (Frankfurt, Germany), RAD51 and ZO1 from Abcam (Cambridge, UK), H3ac and H4ac from Active Motif (Carlsbad, California, USA), p-AMPKα (Thr172), Chk1, p-Chk1 (Ser345), p-GSK3β (Ser9), ac p53 (Lys383), p-p53 (Ser15), p-p70S6K (Thr389) and 53BP1 from cell signaling (Danvers, MA, USA), γH2AX from Millipore (Billerica, MA, USA), p-KAP1 (Ser824) and p-RPA32 (Ser4/Ser8) from Bethyl Laboratories (Montgomery, AL, USA) α-Tubulin ac and β-Actin from Santa Cruz (California, USA), Fluorophore-conjugated secondary antibody Alexa Fluor® 488 and 555 from Life Technologies (Carlsbad, California, USA), anti-rat Alexa Fluor® 488 from Abcam (Cambridge, UK) and peroxidase-conjugated secondary antibody from Rockland (Rockland, Limerick, Pennsylvania, USA). The thymidine analogues 5-Chloro-2-deoxyuridine (CldU) and 5-Iodo-2-deoxyuridine (IdU) were obtained from Sigma (Steinheim, Germany).

### 2.2 Cell culture of mESC and in vitro differentiation

Mouse embryonic stem cells (LF2) were cultivated under feeder-free conditions on 0,1% gelantine-coating using knock-out Dulbecco’s Modified Eagle Medium (KO-DMEM) (gibco, Carlsbad, CA, USA) supplemented with knock-out serum replacement (15%) (gibco, Carlsbad, CA, USA), penicillin/streptomycin (1%), glutamax (1%), β-mercaptoethanol (5 × 10−5 M) (Invitrogen, Carlsbad, CA, USA) and leukemia inhibitory factor (LIF) (Millipore, Billerica, MA, USA) (1000 U/ml) at 37 °C in an atmosphere containing 5% CO_2_. For endothelial differentiation a modified protocol, according to Chiang and Wong [47], was used as described (Fig.1). 0.15×10^5^ cells per 6 well were seeded in serum-free N2b27-medium supplemented with growth factors from day 2 onwards. The differentiation process into EC-like cells were completed on day 6.

**Figure 1:**
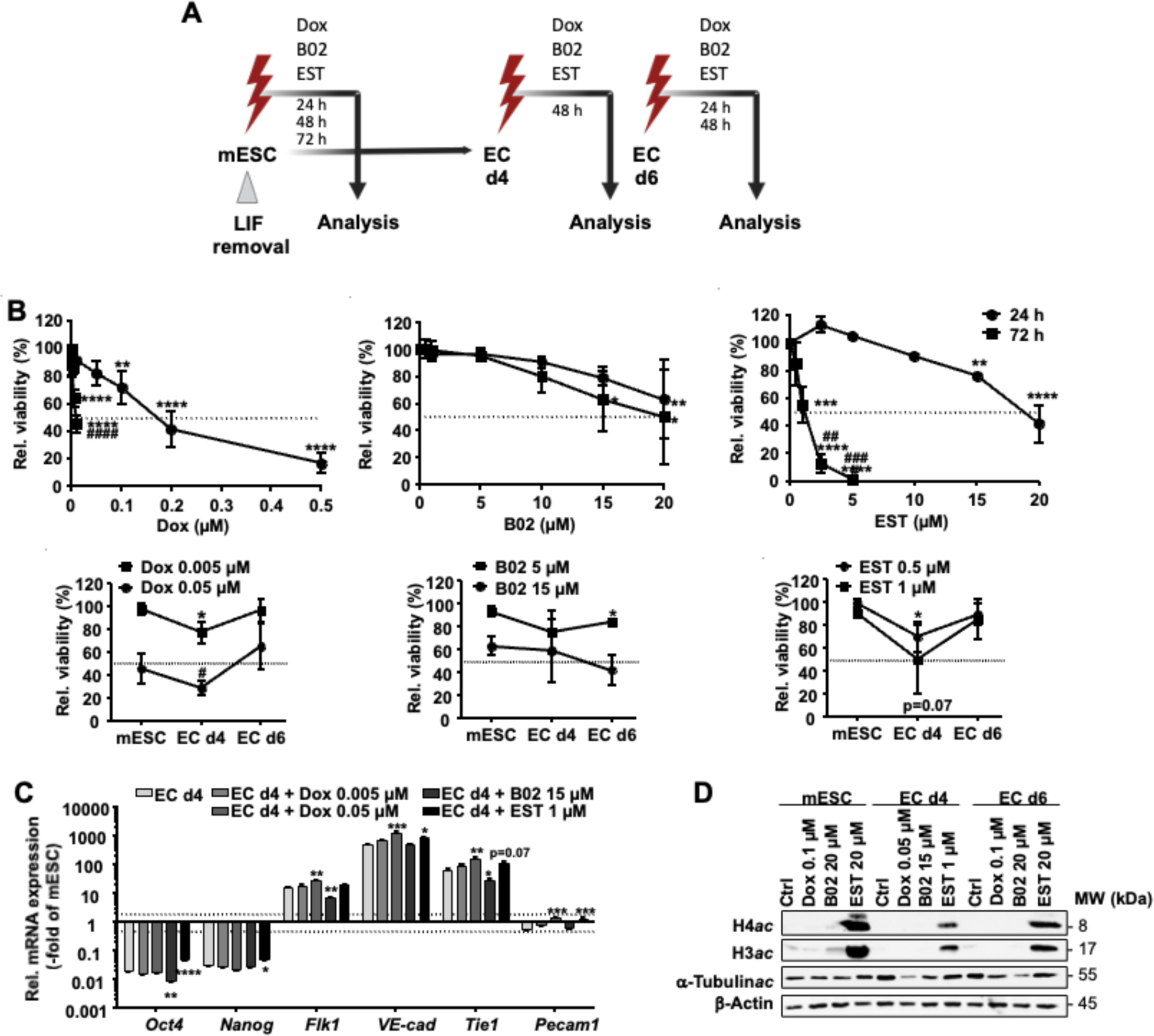
Endothelial progenitor cells (EC d4) reveal higher sensitivity to Dox and epigenetic changes compared to mESC and terminally differentiated EC d6. **A:** Schematic overview of the treatment scheme in course of the differentiation protocol of mESC into EC d6. mESC were treated 24 h after seeding with Dox, B02 and EST for a time period of 24 h, 48 h or 72 h. EC differentiation was induced by removing LIF using serum-free N2b27 medium. On day 2 and day 4 medium was supplemented with growth factors and small molecules. For Analysis cells were treated in progenitor state on day 4 (EC d4) for 48 h continuing differentiation or terminally differentiated on day 6 (EC d6) for 24h or 48 h without growth factor supplement (created with BioRender.com). **B:** Viability of mESC, endothelial progenitor cells (EC d4) and terminally differentiated endothelial-like cells (EC d6) was measured after 24 h or 72 h Dox, B02 and EST treatment using the Alamar Blue assay as described in the methods. Cells were treated for 24 h and 72 h or in course of differentiation for 48 h (EC on day 4, 6 continuing the differentiation protocol, measured on day 6 and 8). Data represent the mean ± SD of three independent experiments performed in quadruplicates. The dashed lines show the IC_50_. One-way ANOVA: p ≤ 0.05*, p ≤ 0.01**, p ≤ 0.001***, p ≤ 0.0001**** as compared to untreated control, Student‘s t–test: p ≤ 0.05^#^ 72 h vs. 24 h. **C:** RT-qPCR gene expression analysis of prototypical stem cell (*Nanog, Oct4*), mesodermal progenitor cell (*Flk1*) and endothelial cell marker genes (*VE-cad, Pecam1, Tie1*). Cells were treated with Dox, B02 and EST on day 4 for 48 h continuing the differentiation protocol. mRNA expression was analyzed on day 6 (terminally differentiated EC). Relative mRNA expression of mESC was set to 1.0. Data represent the mean ± SEM of two independent experiments with two pooled sample per condition performed in triplicates. One-way ANOVA: p ≤ 0.05*, p ≤ 0.01**, p ≤ 0.001*** as compared to untreated control. D: Western Blot analyses of the protein expression of acetylated Histone3 and 4 as well as acetylated α-Tubulin of mESC after 24 h, EC d4 after 48 h and EC d6 after 24 h Dox, B02 or EST treatment. Protein expression of β-Actin was used as loading control.

### 2.3 Analysis of cell viability

Cell viability was determined using the AlamarBlue assay, which measures the reduction of the non-fluorescent dye resazurin (Sigma [Steinheim, Germany]) to fluorescent resorufin. Fluorescence was measured in quadruplicates (excitation: 535 nm, emission: 590 nm, 5 flashes, integration time: 20 μs (Tecan infinite 200, Tecan, Männedorf, Switzerland)) after 2.5-3.5 h incubation with resazurin solution (40µM). Relative cell viability of the corresponding untreated control was set to 100%.

### 2.4 Analysis of cell proliferation

The incorporation of EdU in S-phase was determined using the EdU-Click 488 assay according to the manufacturer’s protocol. Briefly, cells were pulse-treated with 5-ethynyl-2′-deoxyuridine (EdU) for 3 h. Alexa Fluor 488-conjugated dye reaction cocktail was added for 30 min at RT and nuclei were counterstained with DAPI. The percentage of area EdU-positive/area DAPI-positive nuclei was calculated via fluorescence microscopy (Olympus BX43, 20x objective)

### 2.5 Quantitative gene expression analyses (RT-qPCR)

Total RNA was purified using the RNeasy Mini Kit (Qiagen, Hilden, Germany). Reverse transcriptase (RT) reaction was performed with the High Capacity cDNA Reverse Transcription Kit (Applied Biosystems, Darmstadt, Germany) using 900-2000 ng of RNA. Quantitative RT-qPCR analysis was performed using following conditions: 1. 95 °C – 10 min; 2. 45 amplification cycles with 95 °C – 15 s, 55 °C – 15 s, and 72 °C – 17 s; 3. 95 °C – 1 min, 55°C – 1 min, 65°C – 5 s. If not states otherwise, analyses were performed in technical duplicates (array analysis triplicates) using pooled samples from biological duplicates of n=2 independent experiments by use of a CFX96 cycler (BioRad) and the SensiMix SYBR Kit (Bioline, London, UK). The primers used for mRNA expression analyses are listed in Supplementary table 1. To ensure the specificity of the amplification product, melting curves were recorded at the end of the run. mRNA expression levels were normalized to that of *β-Actin* and *Gapdh*. Having checked the target stability value of *Gapdh* and *β-Actin* after each run of the RT-qPCR, we found that the coefficient variance and the M value of both genes were homogeneous in mESC and EC, making these genes useful as housekeeping genes. Relative mRNA expression in mESC or corresponding untreated controls was set to 1.0. Only changes in gene expression of ≤ 0.5- and ≥ 2-fold were considered as biologically relevant.

### 2.6 Western blot analysis

Total protein cell extracts were obtained upon sonication in RIPA buffer (EpiShearTM Probe sonicator, Active Motif [La Hulpe, Belgium]). Protein concentrations were determined using the DC Protein Assay (BioRad [Munich, Germany]). Extracts were supplemented with Roti®-Load buffer (Carl Roth GmbH [Karlsruhe, Germany]) followed by heating (95°C for 5 min). 20 µg of isolated protein was separated by SDS-PAGE and transferred onto a nitrocellulose membrane. The membrane was blocked (5 % non-fat milk in TBS/0.1 % Tween 20 or 5 % BSA in TBS/0.1 % Tween 20) for 1 h at RT and incubated with the corresponding primary antibodies (1:500-5000) overnight at 4°C. After washing the membrane with TBS/0.1 % Tween 20, incubation with peroxidase-conjugated secondary antibody (1:2000) was performed (2 h, RT). The ChemiDoxTM Touch imaging system (BioRad, Munich, Germany) was used for visualization.

### 2.7 Immunofluorescence analysis

To examine the formation of intercellular junctions, the expression of the intracellular tight junction protein ZO1 was analyzed. After fixation (3,7 % formaldehyde/PBS; 15 min, RT), permeabilization (0.1 % TritonX/PBS, 10 min, RT) and blockage (5 % BSA/PBS; 1 h, RT) cells were incubate with primary antibody directed against ZO1 (1:100) at 4 °C overnight. After washing with PBS fluorescence-labeled secondary antibody (Alexa Fluor 488 goat polyclonal to rabbit or rat) was applied (1:1000; 2h, RT). Nuclei were counterstained with DAPI. Microscopic analysis was performed with an Olympus BX43 fluorescence microscopy (100x objective)

### 2.8 Analysis of DNA damage formation and DNA repair

The formation of DNA double-strand breaks (nuclear γH2AX foci) and DNA DSB repair via NHEJ (nuclear 53BP1 foci) were monitored. To this end, cells were fixed with 3.7 % formaldehyde/PBS (15 min, RT) and permeabilized (0.3 % TritonX/PBS; 15 min, RT). After blocking in 5 % BSA in 0.3 % TritonX/PBS (1 h, RT) incubation with γH2AX antibody (1:1000) and 53BP1 antibody (1:500) was done overnight at 4 °C followed by washing with PBS and incubation with fluorescence-labeled secondary antibody (1:500) for 2 h at RT. Nuclei were counterstained with DAPI and were analyzed via fluorescence microscopy (Olympus BX43, 100x objective). In addition, the formation of DNA strand breaks and apurinic/apyrimidinic sites was analyzed via the alkaline comet assay [34]. Cell suspension was mixed with 0.5 % low melting point agarose and applied onto glass slides coated with 1.5 % agarose. After lysis in alkaline buffer (pH 10, 1 h, 4 °C, protected from light) and DNA denaturation in precooled electrophoresis buffer (pH > 13, 25 min, 4 °C, protected from light) electrophoresis was performed (300 mA, 25 V, 4 °C, 25 min). After neuralization DNA was stained with propidium iodide (50 µg/mL) and comets were analyzed via fluorescence microscopy. Quantification of migrated DNA was performed with the TriTek Comet ScoreTM software (version 1.5), evaluating 50 cells per experimental condition.

### 2.9 Analysis of doxorubicin transport

The active import and export of Dox was examined via flow cytometry. Cells were pulse-treated with 0.25 µM and 1 µM Dox for 2 h (import) followed by a 6 h post-incubation with fresh medium (export). After trypsinization, cells were washed twice with cold PBS and pelleted by centrifugation (200 x g, 5 min, 4 °C). After an additional washing step cells were resuspended in PBS. Quantification of Dox import/export was done by flow cytometry-based analyses (excitation: 488 nm, filter: 585/40; Becton Dickinson, AccuriTM C6 plus (Heidelberg, Germany)). Import was defined as an increase in mean intensity and export as a decrease in mean intensity.

### 2.10 DNA fiber spreading assay

For analysis of DNA replication fork progression, the DNA fiber spreading assay was performed as described [35, 36]. Briefly, cells were pulse-labeled with 20 µM CldU (20 min, 37 °C, 5 % C0_2_) and immediately afterwards with 200 µM IdU (20 min, 37 °C, 5 % C02). Cell suspension was applied onto glass slides followed by addition of lysis buffer (0.5% SDS, 200 mM Tris–HCl, 50 mM EDTA). After 6 min incubation (RT), slide were tilted upwards to stretch the fibers, dried 6 min lying horizontally flat and fixed 5 min (RT) in methanol:acetic acid (3:1). After drying (7 min, lying horizontally flat) slides were stored overnight in 70 % ethanol (4 °C). The next day samples were incubated in 100 % methanol (5 min, RT) followed by denaturation in 2.5 M HCl (1 h, RT) and blocking (5 % BSA in PBS, 1 h, 37 °C). DNA Fibers were stained with rat anti-BrdU (1:40) for CldU detection and mouse anti-BrdU (1:70) for IdU detection (in 0.5 % BSA/PBS; 1 h, RT). After washing (0.05% Tween 20 in PBS and PBS) incubation with secondary antibodies anti-rat 488 (1:400) and anti-mouse 555 (1:250) was performed (1 h, RT, 0.5% BSA in PBS). Following washing with 0.05 % Tween 20 in PBS and PBS, the slides were mounted with Fluoroshield (Sigma [Steinheim, Germany]). DNA fibers were analyzed microscopically (Olympus BX43 fluorescence microscope [40 objective]). The length of CldU (green) and IdU (red) tract was measured using the Image J software, analyzing 200 fibers per experimental condition.

### 2.11 Mitochondrial analysis - Mitostaining

Mitochondrial membrane potential (ΔΨ_m_) was analyzed via a live-cell imaging applying a TMRM-MitoTracker Green co-staining approach. To this end, cells were seeded and differentiated on Poly-D-Lysine-fibronectine-coated live-imaging dishes. For the staining, the cells were incubated with 200nM MitoTracker Green (Invitrogen) and 50 nM TMRM (Invitrogen) in DMEM (30 min, 37 °C, 5 % CO_2_). After washing twice with PBS live-cell microscopy was performed using a spinning disc confocal microscope (PerkinElmer) equipped with a 60x oil-immersion objective (N.A = 1.49) and a Hamamatsu C9100 camera (1,000 × 1,000 pixel) (excitation (MitoTracker Green): 488 nm, excitation (TMRM): 561 nm). The cells were kept at 37 °C in OptiMEM supplemented with 10 mM HEPES for the imaging duration. The images were obtained at emission wavelength of 527 nm (W55) and 615 nm (W70) for 488 nm and 561 nm excitation, respectively. The mean intensity of TMRM was measured using the Image J software, analyzing 50 mitochondria per experimental condition (EC d4 + Dox 0.05, EC d6 + Dox 0.1 µM: ≥ 20 mitochondria).

### 2.12 Functional analysis of EC

To monitor prototypical functions of endothelial cells, the uptake of fluorescent 1,1-dioctadecyl-3,3,3,3-tetra-methylindocarbocyanine-labeled acetylated LDL (Dil-acLDL) was examined according to the manufacturer’s protocol. Briefly, cells were incubated 10 μg/ml of Dil-acLDL was added to the culture medium. After 4 h incubation at 37 °C, cells were fixed with 3,7% cold formaldehyde/PBS (15 min, RT) and nuclei were counterstained with DAPI. The uptake of DiI-acLDL was analyzed by fluorescence microscopy (Olympus BX43, 20x objective). As additional surrogate marker of endothelial functionality, their response to inflammatory cytokines was measured. Cells were treated with a pro-inflammatory cytokine mixture (TNFα/IL-1β, 10 ng/mL) for 1 h followed by mRNA expression analysis of activated EC markers *E-selectin, Icam-1, Vcam-1, Ccl2, eNos, iNos* via RT-qPCR.

### 2.13 Statistical analysis

For statistical analysis, the two-tailed unpaired Student’s t-test and One-way ANOVA with Tukeýs and Dunnett’s post-hoc test were employed (using GraphPad Prism 10 software). P-values ≤ 0.05 were considered as statistically significant.

## Results and discussion

### 1. Drug treatment influences the viability of mESC, EC d4 and EC d5 as well as the differentiation potential of mESC in an agent and cell-type specific manner

To comparatively analyze the Dox response of mESC and thereof derived differentiating and terminally differentiated progeny, cell viability was analyzed up to 72 h after drug addition **(Fig. 1A).** For control, pharmacological inhibitors of DSB repair by homologous recombination (HR) and DDR-related factors were included. Apart from the B02, which inhibits the HR-related DSB repair protein RAD51 [37] and is also involved in the regulation of replicative stress responses [38, 39], the HDAC class I inhibitor entinostat (EST) was employed because HDACi are known to interfere with both mechanism of DSB repair and DDR [40, 41].

Data obtained show that 72 h of Dox treatment of mESC causes a massive loss of cell viability as compared to a 24 h treatment period (IC_50_ 24 h: 0.17 µM; IC_50_ 72 h: 0.009 µM) **(Fig. 1B)**. Identical effects were observed upon use of the HDACi EST (IC_50_ 24h: 18.8 µM; IC_50_ 72 h: 2.6 µM), whereas cytotoxicity evoked by B02 was very similar after a treatment period of either 24 h or 72 h (IC_50_ ∼20 µM) **(Fig. 1B)**. This data demonstrates cross-sensitivity of mESC to the genotoxic anticancer drug Dox and the HDACi EST, indicating that both compounds may inhibit cell viability of mESC by overlapping molecular mechanism. To address the question whether cellular drug susceptibility changes during the differentiation process, mESC, thereof derived EC d4 as well as terminally differentiated EC d6 were exposed to Dox and DDR-related pharmacological inhibitors and viability was analyzed 48 h after drug administration **(Fig. 1B)**. The data obtained demonstrate a significant higher Dox and EST susceptibility of EC d4 as compared to both mESC and EC d6 **(Fig. 1B)**, pointing to transient particularly drug sensitive time window during differentiation. This difference was not observed for the RAD51 inhibitor B02 **(Fig. 1B)**. Under our experimental setting of permanent drug treatment, the drug sensitivity of mESC and EC d6 was similar **(Fig. 1B)**. Previously, different Dox sensitivity has been reported between mESC and EC d6 upon short-time Dox pulse-treatment period followed by a post-incubation period of 48 h in the absence of the drug [42]. Thus, differentiation dependent alterations in drug sensitivity clearly depend on the respective treatment conditions, with high-dose short-time treatment regimen giving different results as compared to low-dose long-term treatment plans. This observation is highly important for a meaningful hazard assessment in drug testing aiming to predict developmental toxicity.

In view of the transient Dox and EST hypersensitivity of differentiating EC d4, we ask the question whether drug exposure of differentiating EC d4 impacts their differentiation capacity and correctness of differentiation into EC d6. To answer this question, EC d4 were treated during the remaining differentiation period (i.e. 48 h) with Dox, EST or B02 and the mRNA expression of stem cell factors (*Oct4, Nanog*) and prototypical endothelial factors (*Flk1, VE-cadherin, Tie1, Pecam1*) was analyzed at the end of the differentiation process (i.e. day 6) **(Fig. 1C)**. Data obtained from these analyses reveal moderate, yet statistically significant, differences between the various experimental groups. Dox treatment of EC d4 did not impact the decrease in the mRNA expression of prototypical stem cell factors, which is usually going along with the differentiation process, in the terminally differentiated EC d6. This indicates that Dox induced damage of EC d4 does not impair their differentiation capacity on the level of downregulated mRNA expression of stem cell factors. Importantly, Dox exposure of EC d4 resulted in an upregulated mRNA expression of EC-related marker genes *Flk1, VE-cadherin, Tie1* and *Pecam1* as compared to the non-treated endothelial progenitor control **(Fig. 1C),** indicating that Dox may promote the differentiation of endothelial progenitors. Apparently, Dox treatment of EC d4 differently influences the expression of stem cell- and differentiation-related marker genes. The outcome of B02 and EST treatment was different from that of Dox exposure. EST treatment of endothelial progenitor cells (EC d4) mitigated the differentiation dependent decrease in *Oct4* and *Nanog* mRNA expression and increased the mRNA expression of *VE-cadherin* in terminally differentiated EC d6. By contrast, treatment of EC d4 with B02 promoted the differentiation dependent reduction in *Oct4* mRNA expression while inhibiting the mRNA expression of *Flk1* and *Tie1* in EC d6 **(Fig. 1C).** Overall, the data demonstrate that drug exposure of endothelial progenitor cells during their differentiation process significantly impacts the mRNA expression of differentiation-related maker genes in the terminally differentiated progenitors in an agent-specific manner.

In view of the complex epigenetic alterations going along with the differentiation process, we exemplarily monitored the acetylation status of nuclear histone proteins (H3, H4) and the cytosolic protein α−Tubulin by Western blot. Basal protein levels of H3ac and H4ac were very low in both mESC, EC d4 and EC d6 and specifically increased in response to EST treatment only **(Fig. 1D)**. Notably, this EST-triggered response was most profound in mESC and weakest in EC d4 cells. The acetylation of cytosolic α-tubulin was not influenced in mESC by any of the drugs but was reduced upon treatment with Dox and B02 in EC d4 and EC d6, respectively **(Fig. 1D)**. This data indicates that Dox selectively impacts the acetylation status of cytosolic α-Tubulin but not histone proteins and, moreover, that the responsiveness of mESC, EC d4 and EC d6 to HDAC class I inhibitor EST and the RAD51 B02 inhibitor fluctuates in the course of the differentiation process in a cell-type and agent specific manner.

### 2. Differentiation dependent alterations in drug transport

Since Dox is subject of active transport mechanisms, we investigated whether mESC, EC d4 and EC d6 differ in their Dox import and export capacity. Exploiting the inherent reddish fluorescence of doxorubicin, drug uptake into the cells was measured after a 2h Dox pulse-treatment period (0.25 µM, 1.0 µM). We found a dose-, time- and differentiation status dependent increase in the intracellular concentration of Dox. At a low Dox concentration (i.e. 0.25 µM), a significantly increased drug import was only detected in EC d4 **(Fig. 2A).** Following treatment with higher Dox concentration (1 µM), further increase in drug import was found for all three cell types with EC d4 and EC d6 displaying a significantly higher maximum intracellular fluorescence intensity as compared to mESC **(Fig. 2A).** Hence, Dox import capacity is particularly high in EC d4 and increases in the course of the endothelial differentiation process. To monitor the cellś drug export capacity, residual intracellular Dox fluorescence was measured after a subsequent 6 h post-incubation period in fresh medium in the absence of Dox. For all three cell types, the kinetic of drug export was found to depend on the loading dose initially used for the pulse treatment **(Fig. 2A)**. EC d6 show a tendentially increased residual intracellular Dox level as compared to mESC. Calculation of the Dox efflux capacity revealed an enhanced export activity of EC d4 as compared to the other differentiation stages **(Fig. 2B).** Thus, following an efficient drug uptake of Dox, mechanisms are triggered in EC d4 which in turn lead to an effective removal of Dox from these cells.

**Figure 2:**
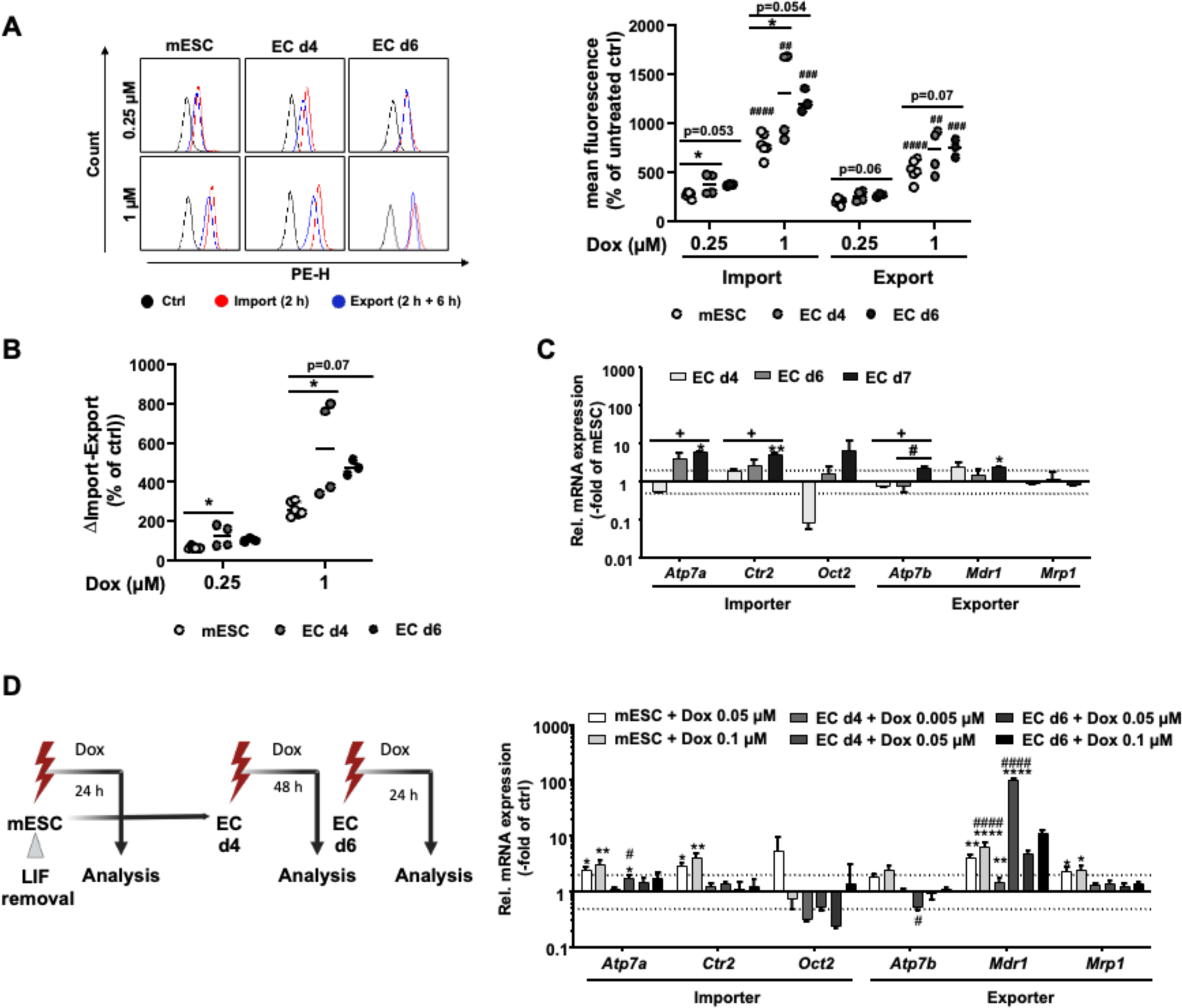
Differentiation dependent alterations in drug transport. **A:** Intracellular doxorubicin (Dox) fluorescence was measured via flow cytometry and was taken indicative for Dox import and export. mESC, EC d4 and EC d6 were treated for 2 h with 0.25 µM and 1 µM Dox. To analyze Dox import increase in mean fluorescence intensity was measured via flow cytometry immediately after pulse treatment or after a 6 h post incubation period (export). **B:** To calculate the export capacity the Dox efflux normalized to the corresponding untreated control was calculated. Data represent three to four independent experiments. One-way ANOVA as compared to mESC (p≤0.05*), Student‘s t–test: 1 µM vs. 0.25 µM (p ≤ 0.01^##^, p ≤ 0.001^###^, p ≤ 0.0001^####^) **C:** Basal mRNA expression of selected drug importer (*Atp7a, Ctr2, Oct2*) and exporter (*Atp7b, Mdr1, Mrp1*) genes of EC d4 and EC d6 and 7 in relation to mESC. Data show the mean ± SEM of two to three independent experiments with two pooled samples each performed in duplicates. Changes in mRNA levels of ≥ 2.0 and ≤ 0.5 indicated by dashed lines are considered as biologically relevant. One-way ANOVA: p ≤ 0.05*, p ≤ 0.01** as compared to mESC, + EC d4 vs, EC d7, # EC d6 vs d7. **D:** Schematic overview of the drug-treatment scheme (left panel; created with BioRender.com). Right panel: Comparative RT qPCR analysis of mRNA expression of selected drug transporter genes of mESC, EC d4 and EC d6 after Dox exposure. mESC and EC d6 were treated for 24 h or EC d4 for 48 h continuing differentiation protocol with 1-2 equitoxic doses of Dox. Relative mRNA expression of the corresponding untreated control was set to 1.0. Data show the mean ± SEM of two to three independent experiments with two pooled samples per condition each performed in duplicates. Changes in mRNA levels of ≥ 2.0 and ≤ 0.5 indicated by dashed lines are considered as biologically relevant. One-way ANOVA: p ≤ 0.05*, p ≤ 0.01** as compared to mESC, + EC d4 vs. EC d7, # EC d6 vs d7

To identify the transporters involved, mRNA expression of selected Dox importers and exporters was investigated under basal situation and after Dox treatment. Data obtained show differences in the basal mRNA expression of drug importers, notably the importers *Atp7a* and *Oct2* and the exporter *Atp7b,* between mESC, EC d4 and terminally differentiated EC d6 (**Fig. 2C)**. Following Dox treatment, the mRNA expression of the drug importers *Atp7a*, *Ctr2* and *Oct2* was specifically enhanced in mESC **(Fig. 2D)**. Among the drug exporters, the most substantial Dox-stimulated increase was observed for *Mdr1*, with EC d4 showing the strongest response at an equimolar Dox concentration of 0.05 µM **(Fig. 2D)**. This finding indicates that the elevated Dox export activity of EC d4 **(see Fig. 2B)** is likely due to an enhanced MDR1-related drug export activity. Overall, we observed differentiation dependent changes in Dox transport capacity, both under basal situation and following Dox treatment, with differentiating EC d4 revealing the most pronounced differences in the mRNA expression of drug transporters as compared to mESC and EC d6. Yet, having in mind that EC d4 cells reveal a significantly higher Dox sensitivity than mESC and EC d6, the observed alterations in drug transporter expression do not provide a rationally convincing molecular explanation for their Dox hypersensitivity. Hence, we conclude that the Dox hypersensitive phenotype of EC d4 is likely independent of differentiation-dependent alterations in the expression of transport-related pre-target mechanisms that can contribute to drug resistance.

### 3. Differentiation dependent formation of DNA damage by doxorubicin

Inhibition of Topo II by Dox is known to cause the formation of DSB, which are highly cytotoxic DNA lesions and potent triggers of the DDR [3]. They can be repaired by homologous recombination (HR) or non-homologous end-joining (NHEJ) [43, 44]. The steady state level of DSB by Dox and pharmacological DDR/repair inhibitors was comparatively investigated in mESC, EC d4 and EC d6 by monitoring the number of nuclear γH2AX and 53BP1 foci, both of which are well accepted surrogate markers of DSB [45–48] **(Fig. 3B-D)**. For these analyses, equitoxic concentrations of the various drugs were used for treatment **(Fig. 3A)**. A significant increase in the number of nuclear γH2AX foci was observed in mESC following treatment with Dox and the Topo II inhibitor etoposide (Eto), which was included for control, as well as with B02 and EST **(Fig. 3B)**. Similar results were obtained measuring the formation of nuclear 53BP1 foci **(Fig 3B)**, which are considered as marker of DSB that are processed by NHEJ [44, 49]. Similarly, EC d4 also responded to treatment with Dox, Eto, B02 or EST with a clear increase in DSB **(Fig. 3C)**. As opposed to mESC, EC d6 only revealed a significant increase in the number of DSB upon Eto and Dox exposure, but not following treatment with the aforementioned DDR/repair inhibitors **(Fig. 3D)**. Moreover, the steady state number of Dox-induced γH2AX foci was highest in EC d6 **(Fig. 3B-D)**. Comparing the number of nuclear 53BP1 foci in cells of various differentiation stages, highest number was again found in EC d6 **(Fig. 3B-D).** Overall, the data support the hypothesis that the capacity of DSB repair by HR declines with the differentiation process, while NHEJ increases [50]. Measuring the formation of SSB by use of the alkaline comet assay, a significant increase in SSB was observed in mESC treated with Eto, Dox or EST **(Fig. 3E),** while no or only minor increase in the steady-state level of DNA single strand breaks (SSB) was found in drug-treated EC d4 and EC d6, respectively **(Fig. 3F and 3G)**. This finding indicates that differentiating and terminally differentiated cells are more resistant to the formation of SSB by the drugs used, or, alternatively, repair SSB more efficiently than mESC. Here, base excision repair (BER) may be a candidate repair pathway, which is of particular relevance for SSB repair and, moreover, has been shown to change during differentiation [51–53].

**Figure 3:**
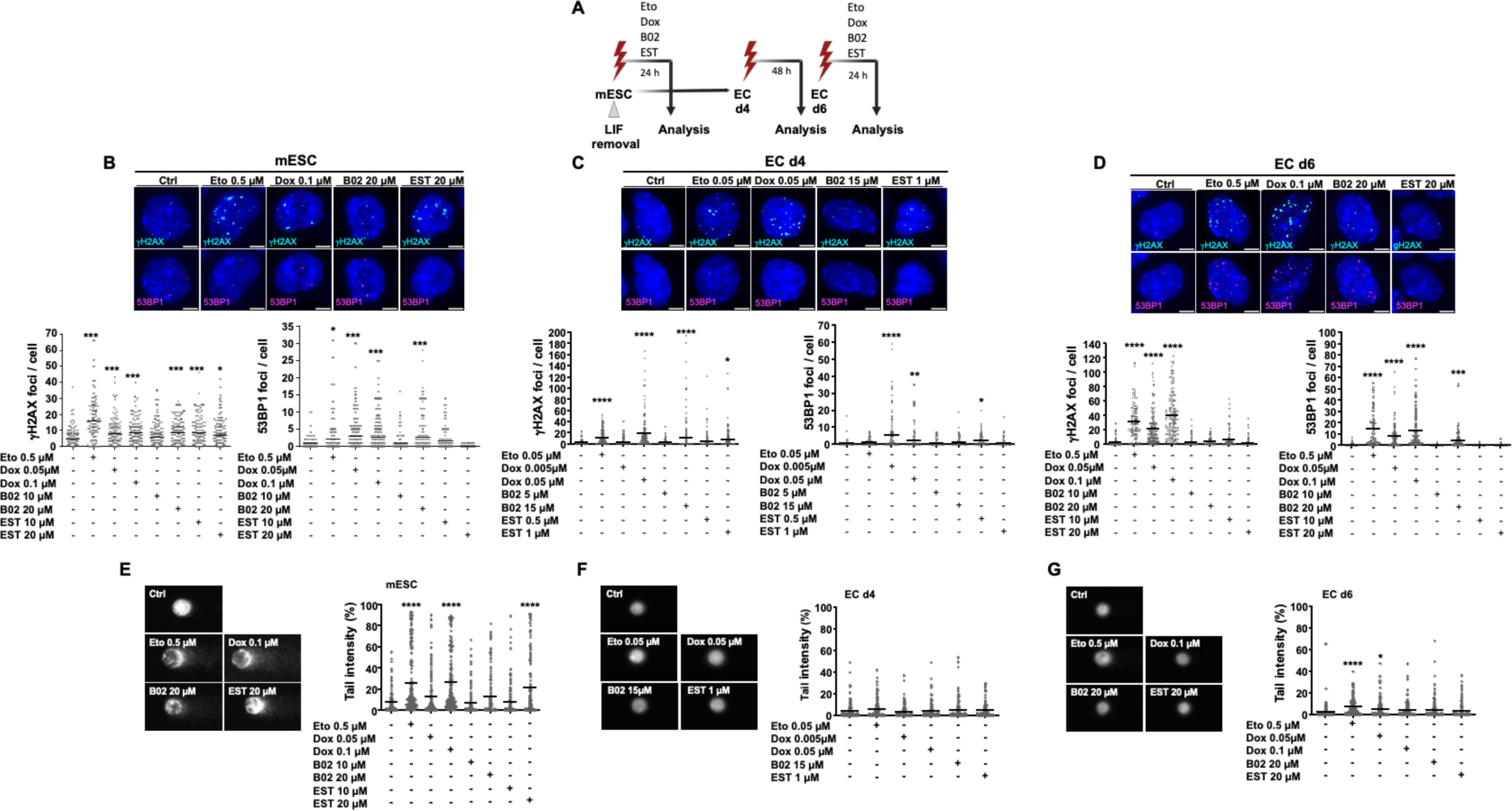
The type of steady state DNA damage is agent- and cell type-specific. Cells were treated undifferentiated (mESC) and terminally differentiated (EC d6) on day 6 for 24 h or on day 4 (EC d4) for 48 h continuing differentiation protocol with 1-2 equitoxic doses of Dox, EST and B02. As an additional control, cells were treated with one dose of TopoII inhibitor Etoposide (Eto). **A:** Schematic overview of the differentiation protocol of mESC into EC and drug-treatment scheme in the course of the differentiation process (created with BioRender.com). **B-D:** Results of immunocytochemical analyses of nuclear γH2AX (cyan) and nuclear 53BP1 (magenta) foci formation per cell (Dox, B02, EST: n=3, N=50; Eto: n=2, N=50). in mESC (B), EC d4 (C) and EC d6 (D) following drug treatment. One-way ANOVA: p ≤ 0.05*, p ≤ 0.01**, p ≤ 0.001***, p ≤ 0.0001**** as compared to the corresponding untreated control. Scale bar: 5 µm. **E-G:** The steady state levels of DNA single strand breaks (SSB), DNA double-strand breaks (DSB) in mESC (E), EC d4 (F) and EC d6 (G) after treatment was analyzed by use of the alkaline comet assay. Quantification of DNA in tail (%) was performed as described in the methods (n=3, N=50). One-way ANOVA: p ≤ 0.05*, p ≤ 0.0001**** as compared to the corresponding untreated control.

### 4. Doxo and pharmacological inhibitors differently impact replicative stress responses of mESC, EC d4 and EC d6

To monitor alterations in proliferation going along with differentiation and/or drug treatment, S-phase activity of mESC, EC d4 and EC d6 was monitored by measuring the incorporation of EdU under basal situation and following drug treatment. Under basal situation, the percentage of EdU positive cells dropped with the differentiation status from 70 % EdU positive mESC to 45-30 % EdU positive EC d4 and EC d6 **(Fig. 4B-D)**. Dox treatment caused a significant drop in the number of EdU positive cells in mESC **(Fig. 4B)**. Similar effect was found upon treatment of mESC with EST but not B02 **(Fig. 4B)**. EC d4 respond to treatment with a high Dox concentration and following B02 treatment with a decrease in S-phase activity, while this effect was not observed after EST exposure **(Fig. 4C)**. Noteworthy, the number of EdU positive EC d6 cells remained unaffected by drug treatment. In summary, this data show agent- and differentiation-dependent alterations in the S-phase activity following drug-induced insults.

**Figure 4:**
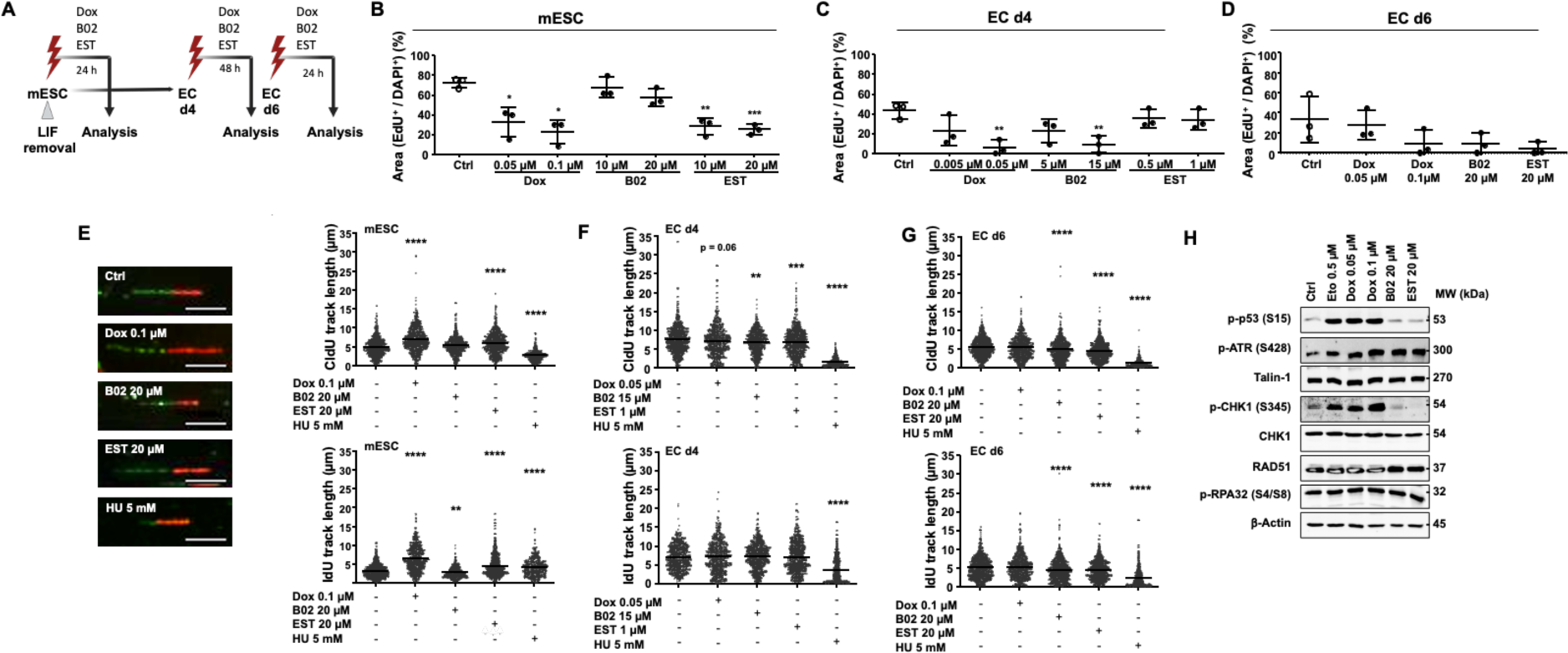
Influence of Dox, B02 and EST treatment on proliferation and activation of replication stress-associated mechanisms. Cells were treated undifferentiated and terminally differentiated on day 6 for 24 h or on day 4 for 48 h continuing differentiation protocol after treatment with the indicated concentrations of Dox, EST or B02. **A:** Schematic illustration of the treatment scheme (created with BioRender.com). **B-D:** Cell proliferation was measured after drug treatment by monitoring EdU incorporation (3 h EdU pulse-treatment) The percentage of EdU^+^ cells was calculated by analyzing the EdU^+^ area in correlation to the DAPI counterstain (fluorescence microscopy, 20x objective). Data represent the mean ± SD of three independent experiments. One-way ANOVA: p ≤ 0.05*, p ≤ 0.01**, p ≤ 0.001*** as compared to the corresponding untreated control. **E-G:** Replication fork progression was analyzed by use of the DNA fiber spreading assay as described in methods. Data shown were obtained from n=3-4 independent experiments with 200 fibers being analyzed per experimental sample. For additional control, cells were treated with the ribonucleotide reductase inhibitor hydroxy-urea (HU; 5 mM, 1 h) (n=2, N=1-2). One-way ANOVA: p ≤ 0.01**, p ≤ 0.001***, p ≤ 0.0001**** as compared to the corresponding untreated control. E (left panel): Representative images of immunocytochemical analyses of CldU(green)/IdU (red) containing DNA fibers in mESC (Scale bar 5 µm). **H:** Protein expression of selected replication stress response-related factors of mESC non-treated and following 24 h Eto, Dox, B02 and EST treatment were analyzed by Western blot. Protein expression of β-Actin and Talin-1 were used as loading controls.

To further clarify the impact of the drugs on the progression of the replication fork in relation to the cellś differentiation status, the DNA fiber spreading assay was employed **(Fig. 4E-G)**. Untreated mESC displayed a reduced replication fork progression as compared to EC d4, as indicated by short-length tracks during both pulses **(Fig. 4E)**. Treatment with the ribonucleotide reductase inhibitor hydroxy-urea (HU) decreases replication fork speed in all cell types under investigation, yet with EC d4 revealing the strongest response **(Fig. 4E-G).** Inhibition of RAD51, which is involved in the regulation of HR and of replicative stress responses [43, 54], did not influence replication fork speed in mESC but revealed inhibitory effects in both EC d4 and EC d6 **(Fig. 4E-G)**. By contrast, interference with mechanisms of the DDR and/or DNA repair by the class I HDACi EST [55] caused increased incorporation of fluorescence-labeled nucleotides in mESC, while having the opposite effects in both EC d4 and EC d6 **(Fig. 4E-G).** Exposure of mESC to Dox also resulted in an accelerated fork progression but with visible gaps within the track **(Fig. 4E)**. By contrast, Dox had no influence on fork speed in EC d4 and EC d6 **(Fig 4F-G).** The data demonstrate remarkable drug-dependent differences on the replication fork activity of mESC versus differentiating EC d4 and terminally differentiated EC d6. This finding might be related to replication specific characteristics of mESC. Asynchronous mESC spend most time in S-phase and less time in G1- and G2-phase of the cell cycle [56]. Therefore, factors involved in DNA replication and replication stress response are highly abundant in mESC, which, in addition, are reported to have a compromised G1/S checkpoint [56]. mESC experience replication stress under basal condition, but at the same time they are thought to use specific mechanisms like fork slowing and fork remodeling to ensure genome maintenance during replication. With ongoing differentiation, basal replicative stress level decreases and cells are spending more time in gap phases for template quality control before entering S-phase [56].

In extension to this data, the activation status of DDR-associated factors was monitored by Western blot analysis. The tumor suppressor p53 acts as a central regulator in cellular stress responses. Therefore, we investigated ATM/ATR-mediated activation of p53 via phosphorylation on Ser15. Both Dox and Eto exposure caused an increase in the protein level of Ser15-phosphorylated p53 in mESC, while these cells did not respond to treatment with B02 or EST **(Fig. 4H)**. Having in mind the key regulatory function of ATR in the regulation of replication-associated stress response [57], the protein level of p-ATR, the ATR-substrate CHK1 (p-CHK1) and ssDNA binding proteins RPA (p-RPA32) and RAD51 were monitored. Dox treatment of mESC increased the level of p-ATR and p-Chk1 **(Fig. 4H)**, which supports the hypothesis of replicative stress evoked by Dox exposure. Noteworthy, B02 and EST increased the protein level of p-ATR but not of p-Chk1 **(Fig. 4H)**. None of the drug treatments caused changes in the protein levels of p-RPA32 or RAD51 **(Fig. 4H)**. Summarizing, the western blot data support the hypothesis of drug-specific stimulatory effects on replication stress-associated mechanisms regulated by the ATR/CHK1/p53 axis.

### 5. Differentiation dependent activation of mechanisms of the DDR by Dox and DDR / DNA repair inhibitors

To figure out whether the expression of Dox target proteins is changing during differentiation, we comparatively analyzed the mRNA expression of the Topo II isoforms in mESC, EC d4 and EC d6. Whereas the mRNA expression of *Top2a* did not change with differentiation, we observed an upregulation of *Top2b* mRNA levels in EC d6 as compared to mESC and EC d4 **(Fig. 5A)**. Since Topoisomerase IIβ is involved in transcriptional regulation [58] and, moreover, is present in mitochondria [59], its upregulation may be related to differentiation dependent alterations in gene expression and/or the metabolic shift from glycolysis towards oxidative phosphorylation (OxPhos) that is known to go along with differentiation [60, 61]. Since Dox can also cause oxidative stress, mRNA expression of antioxidative defense-related factors was monitored. These analyses revealed a downregulation of *Hmox1* and *Nqo1* mRNA level in EC d4 and EC d6 as compared to mESC **(Fig 5B)**. In addition, a selective downregulation of *Nrf2* and *Nos3* mRNA levels was found in EC d4 but not in EC d6 **(Fig. 5B)**, indicating transient changes in the expression of antioxidative functions occurring during endothelial differentiation of mESC. In line with these data, transient fluctuations of antioxidative functions have also been reported during renal differentiation of hiPSC [62].

**Figure 5:**
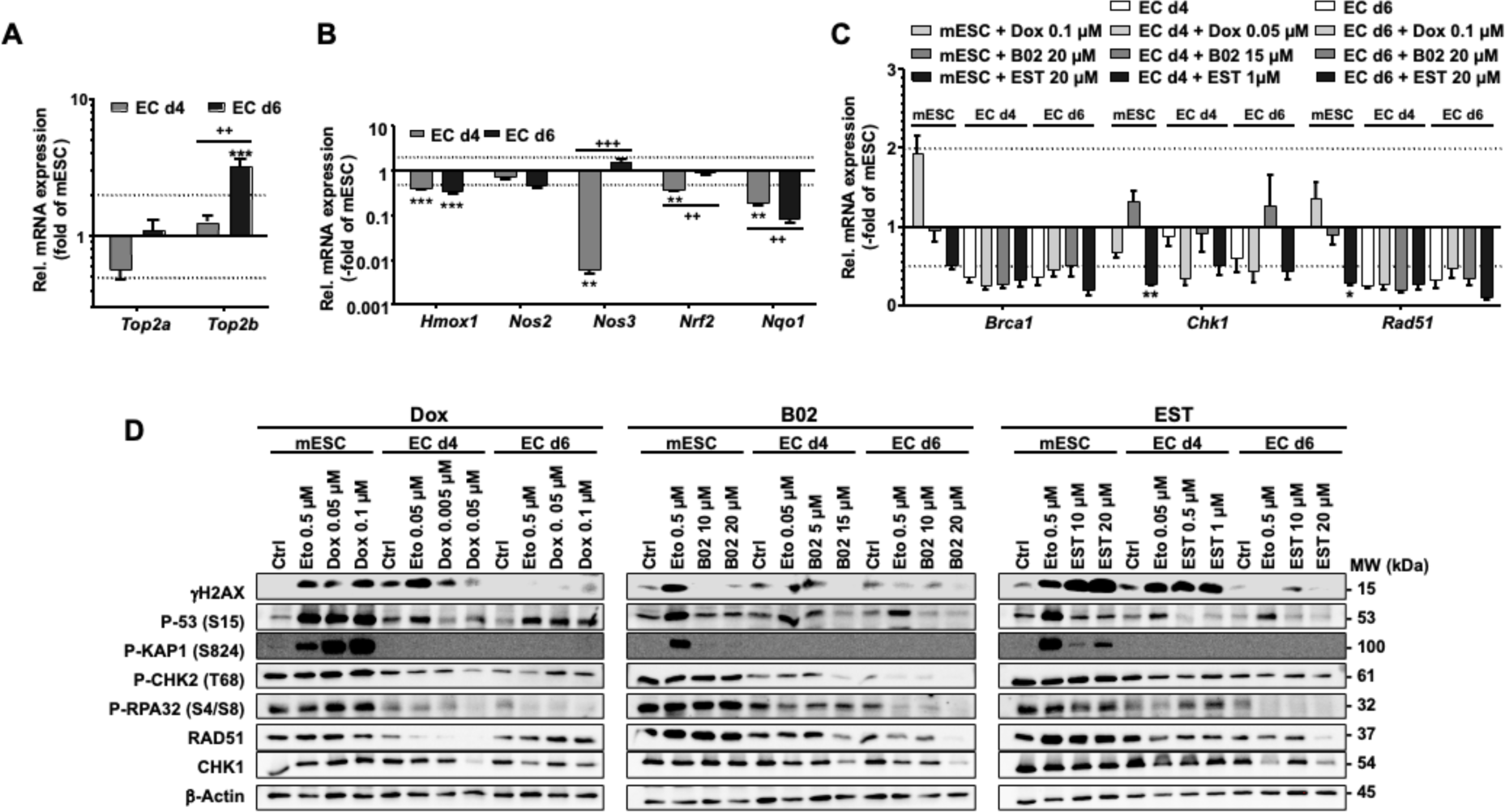
Differentiation dependent activation of mechanisms of the DDR by Dox and DDR/DNA repair inhibitors. **A:** RT-qPCR analysis of basal mRNA expression of *Top2a* and *Top2b* in EC d4 and EC d6 as compared to mESC. Data show the mean ± SEM of two independent experiments each performed in biological duplicates. Changes in mRNA levels of ≥ 2.0 and ≤ 0.5 are indicated by dashed lines and are considered as biologically relevant. One-way ANOVA: p ≤ 0.001*** as compared to mESC, Student’s t-test: p ≤ 0.05+ EC d4 vs EC d6. **B:** RT_qPCR analysis of basal mRNA expression of antioxidative response-related factors in EC d4 and EC d6 as compared to mESC. Data show the mean ± SEM of two independent experiments with two pooled samples each performed in duplicates. Changes in mRNA levels of ≥ 2.0 and ≤ 0.5 are indicated by dashed lines and are considered as biologically relevant. One-way ANOVA: p ≤ 0.01** p ≤ 0.001*** as compared to mESC, Student’s t-test: p ≤ 0.01^++^ p ≤ 0.001^+++^ EC d4 vs. EC d6. **C:** Comparative analysis of mRNA expression of HR- and DDR-associated factors *Brca1, Rad51* and *Chk1*, respectively, following Dox, B02 and EST treatment of mESC, EC d4 and EC d6. mESC and terminally differentiated EC d6 were treated for 24 h, while EC d4 were exposed during the whole differentiation time (i.e. 48 h). Data represent the mean ± SEM of two independent experiments each performed in biological duplicates. Changes in mRNA levels of ≥ 2.0 and ≤ 0.5 are indicated by dashed lines and are considered as biologically relevant. Relative mRNA expression of mESC was set to 1.0. One-way ANOVA p ≤ 0.05*, p ≤ 0.01** as compared to the corresponding untreated control. **D:** Western Blot-based analysis of the protein expression of selected DNA repair and DDR-related factors. mESC and EC d6 cells were treated 24 h and EC d4 for 48 h with each two different doses of Dox, B02 or EST as indicated. As additional positive control, cells were treated with etoposide (Eto). The protein expression of β-Actin was analyzed as loading control.

Since Dox causes the formation of DSB that are subject of DSB repair involving HR and both B02 and EST are known to impair DSB repair and DDR-related factors like RAD51, BRCA1 and CHK1 [40, 63–65], we monitored differentiation dependent alterations in the mRNA **(Fig 5C)** and protein expression **(Fig 5D)** of these factors under basal situation and after drug treatment in mESC, EC d4 and EC d6. mRNA expression of *Brca1*, *Chk1*, *Rad51* remained unaffected or was decreased both under basal situation and following drug treatment in all stages of differentiation **(Fig. 5C).** EST exposure induced a downregulation of *Brca1* in EC d4 and EC d6, while *Chk1* and *Rad51* were found decreased in all cell types **(Fig. 5C)**. Taken together the data show a negative regulation of the mRNA expression of DNA DSB repair related genes, in particular HR-related factors, with ongoing differentiation and upon drug exposure in cells of all differentiation stages. On the protein level, RAD51 and CHK1 were found to be downregulated after high dose of EST in EC d6, but not in the other cell types **(Fig 5D).** This observation reflects the high abundance of replication stress- and homologous recombination-related factors in mESC, which drops with differentiation. Therefore, EST mediated downregulation was only detected at lower level compared to mESC in EC d6.

Analysis of the activation status of DDR-related signaling pathways revealed a strong increase in the phosphorylation of histone H2AX on Serine 139 (γH2AX) in mESC following treatment with the genotoxin Dox and Eto as well as the HDAC_i_ EST, while the Rad51_i_ B02 did not caused an increase in γH2AX levels as compared to the untreated controls **(Fig 5D, middle panel)**. Drug induced increase in the protein levels of p-p53 was only detectable after Dox treatment of mESC and EC d6 but not of EC d4 **(Fig. 5D, left panel)**, indicating that specifically Dox-stimulated activation of p53-related signaling is compromised in differentiating cells. In addition, a substantial Dox-induced increase in the phosphorylation level of the chromatin regulatory factor Kap1 was only observed in mESC but not in EC d4 and differentiated EC d6 **(Fig. 5D).** Noteworthy in this context, cells of all three differentiation stages revealed a similar increase in p-KAP1 levels following treatment with the Topo II inhibitor Eto **(Fig. 5D)**, again highlighting profound genotoxin-specific effects on chromatin-related factors. Neither EST nor B02 simulated an increase in p-KAP1 protein levels in any of the cells under investigation **(Fig. 5D**), demonstrating selective effects of these inhibitors on distinct factors of the DDR.

### 6. Drug treatment of progenitor EC d4 influences mitochondrial membrane potential of differentiated EC d6

Multiple molecular mechanisms contribute to Dox-induced cytotoxicity, including mitochondrial toxicity [66, 67]. Therefore, membrane potential was analyzed via live-cell imaging of MitoTracker^TM^ and TMRM stained cells. The data show that the mitochondria of progenitor EC d4 are affected by all three substances, with 0.05 µM Dox causing the strongest effect, as indicated by a significant drop in mitochondrial membrane potential **(Fig 6A)**. Mitochondrial membrane potential of terminally differentiated EC (EC d6) was also significantly reduced following Dox and B02 treatment, yet remained largely unaffected by treatment with the HDACi EST **(Fig. 6B).** This finding indicate that mitochondrial damage caused by mono-treatment with DNA repair / DDR inhibitors is agent specific. Taken together this data reveal a so far unknown pronounced susceptibility of mitochondria of differentiating cells to substances interfering with genomic stability, including the genotoxin Dox but also pharmacological inhibitors of DNA repair and DDR-related processes. This phenomenon may be related to changes in mitochondrial homeostasis and metabolic activity going along with differentiation [61] and/or mitotoxic effects of genotoxic metabolites that are endogenously generated in varying quantities and/or qualities depending on the cellular differentiation status.

**Figure 6:**
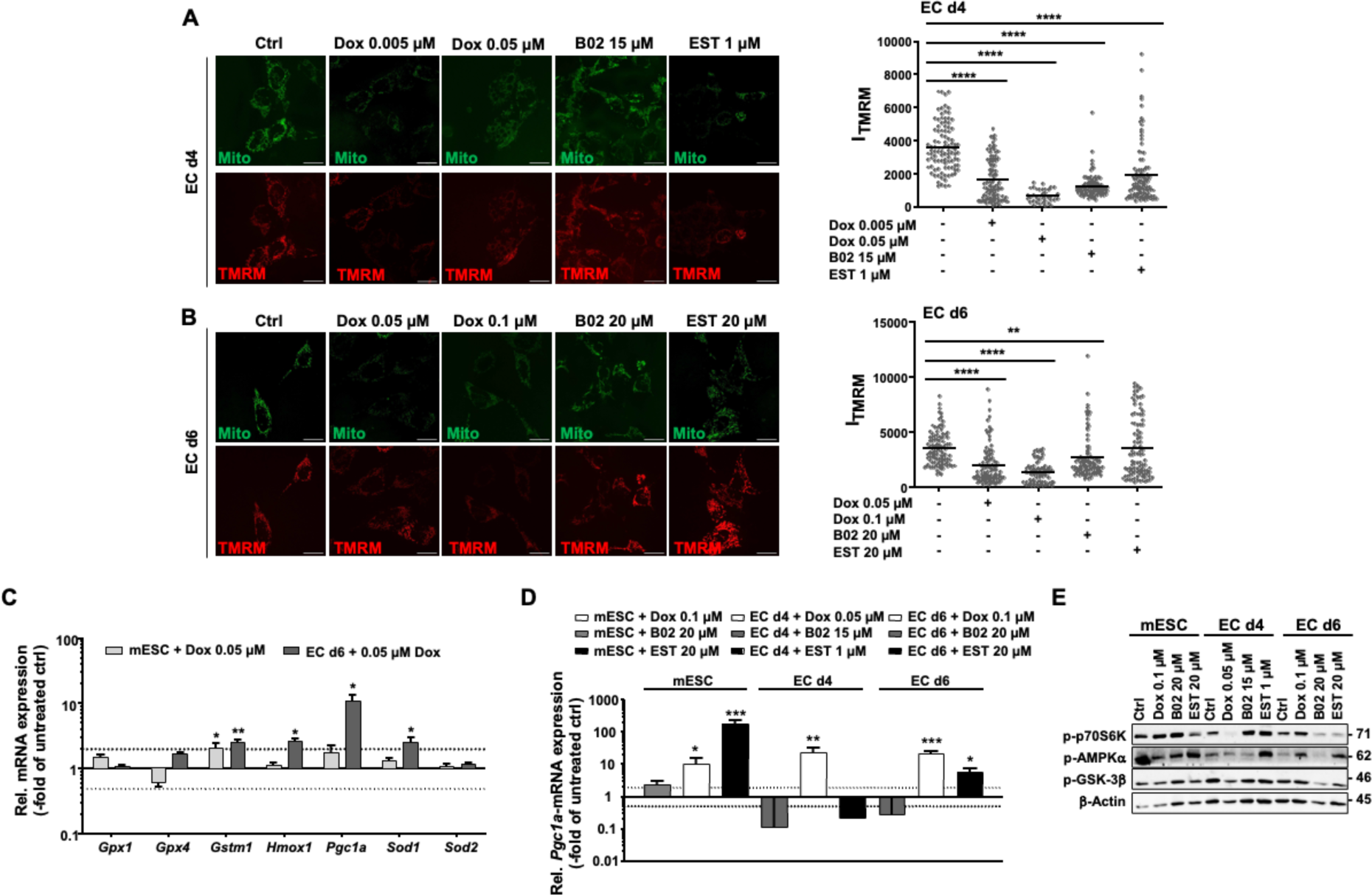
Drug treatment of progenitor EC d4 influences mitochondrial membrane potential of differentiated EC d6. Mitochondrial membrane potential (ΔΨ_m_) after drug treatment was analyzed via a live-cell imaging of a TMRM-MitoTracker^TM^ Green co-staining. **A, B:** Cells were treated on day 4 of the differentiation process (EC d4) for 48 h with Dox, B02 or EST and mitochondrial potential was analyzed in differentiated EC d6 (A). For control, terminally differentiated EC d6 were treated for 24 h with Dox, B02 or EST (B). Following drug treatment, cells were incubated for 30 min with MitoTracker^TM^ Green and TMRM. Green color show mitochondria and red indicates an intact membrane potential. Membrane integrity (fluorescence intensity) was determined as described in the methods. Data show the mean of two independent experiments (N=50, EC d4 + Dox 0.05 µM/ EC d6 + Dox 0.1 µM: N ≥ 20). One-way ANOVA: p ≤ 0.05*, p ≤ 0.01**, p ≤ 0.0001**** as compared to the respective untreated control. Scale bar 20 µm. **C:** Relative mRNA expression of a selected subset of mitochondria-related genes was analyzed in mESC and EC d6 after 24 h Dox exposure (0.05 µM) by RT-qPCR. Relative mRNA expression of genes in the corresponding untreated control was set to 1.0. Data represent the mean ± SEM of triplicate determinations. Only changes in mRNA levels of ≥ 2.0 and ≤ 0.5 were considered as biological relevant (dashed lines). Student’s t-test: p ≤ 0.05*, p ≤ 0.01**, as compared to the respective untreated control. **D:** Comparative RT-qPCR gene expression analysis of *Pgc1* mRNA expression in mESC, EC d4 and EC d6 after treatment with Dox, B02 and EST (mECS, EC d6: 24 h; EC d4: 48 h). Relative mRNA expression of the corresponding controls was set to 1.0. Data represent the mean ± SEM of two independent experiments with two pooled samples each performed in duplicates. Changes in mRNA levels of ≥ 2.0 and ≤ 0.5 indicated by dashed lines are considered as biologically relevant. One-way ANOVA: p ≤ 0.01**, p ≤ 0.001*** as compared to the respective untreated control. **E:** Western Blot analysis of activated (i.e. phosphorylated) protein kinases involved in the regulation of metabolism (p-p70S6K, p-AMPKα and p-GSK-3β) in mESC, EC d4 and EC d6 following Dox, B02 and EST treatment as described above. β-Actin protein expression was used for protein loading control.

Gene expression analysis of a subset of detoxification and mitochondrial function-related genes showed a minor increase in the mRNA expression of the antioxidative factors *Hmox1* and *Sod1* in Dox-treated EC d6 as compared to mESC **(Fig. 6C)**. In addition, a prominent upregulation of *Pgc1a* mRNA levels was observed following Dox treatment of terminally differentiated EC d6 as compared to mESC **(Fig 6C)**. Since PGC1α plays a key role in regulating mitochondrial biogenesis, which is involved in maintaining mitochondrial homeostasis [68], this finding again is in line with the known increase in OXPhos during differentiation. Comparative analysis of Dox-simulated *Pgc1a*-mRNA expression in cells of varying stages of differentiation using higher concentration of Dox revealed an upregulation of this marker gene independent of the cell’s differentiation status **(Fig 6D)**. Treatment with EST also increased *Pgc1a* mRNA levels in mESC and, to a lower extent in EC d6, but not EC d4 **(Fig. 6D),** again pointing to a compromised response of differentiating cells to pharmacological inhibition of HDAC. B02 treatment did not affect *Pgc1a* mRNA expression in any of the cell types under investigation **(Fig. 6D)**. To further investigate whether drug treatment interferes with metabolism, the activation status of the protein kinases p70S6 kinase, AMP kinase alpha and GSK-3β kinase, which play key roles in the regulation of metabolic processes was investigated. The data obtained show profound differentiation- and drug-dependent variations in the activation status of these kinases. For example, the protein level of phosphorylated p70S6K was reduced in EC d4 and EC d6 as compared to mESC under untreated conditions **(Fig. 6E)**, indicating differentiation specific differences in the PI3 kinase pathway and protein synthesis. The same holds true for p-AMPKα, which is important for the regulation of cellular energy homeostasis, while the basal level of p-GSK-3β, which inhibits glycogen synthesis, was slightly enhanced in EC d4 **(Fig. 6E)**. Following drug treatment, the phosphorylation of p70S6K was largely reduced in EC d4 after Dox exposure, while both B02 and EST treatment increased the p-p70S6K level in EC d4. Of note, the opposite was observed in B02 or EST-treated terminally differentiated EC d6 **(Fig 6E**), showing substantial differences in the metabolic response of differentiating versus differentiated cells following treatment with DNA repair / DDR inhibitors. Similarly, multiple agent- and differentiation dependent variations in the the expression level of p-AMPKα and p-GSK-3β were observed, such as increased p-AMPKα levels in EST treated EC d4 and EC d6 cells **(Fig. 6E)**. Taken together, the data demonstrate that mESC, endothelial progenitor EC d4 and terminally differentiated EC d6 cells differently respond to Dox-, B02- or EST-induced damage with either an upregulation or inhibition of multiple mechanisms related to mitochondrial biogenesis, mitochondrial homeostasis and metabolism.

### 7. Impact of drug treatment of differentiating EC d4 on the functionality of differentiated EC d6

Apart from affecting proliferation, triggering cell death or influencing the differentiation process, drug treatment of progenitor cells with subtoxic drug concentrations may also lead to a dysfunction of the differentiated surviving progeny. To scrutinize this hypothesis, EC d4 were treated with Dox or DDR/DNA repair inhibitors followed by the analysis of surrogate markers of prototypical endothelial functions in the differentiated progeny. Against the background of the physiologically highly important barrier function of the endothelium, the expression and localization of the tight junction protein ZO1 was analyzed by immunocytochemistry. Endothelial cells (EC) originating from untreated EC d4 showed homogenous structures of ZO1 containing tight junctions **(Fig. 7A)**. By contrast, EC that were differentiated from Dox-treated EC d4 revealed fragmented / interrupted ZO1 containing tight junctions **(Fig 7A).** Apparently, Dox exposure of progenitor cells impairs the correct formation of tight junctions in the differentiated progeny. Treatment of EC d4 with DDR inhibitors B02 and EST does not affect the presence and assembly of the ZO1 protein into tight junctions **(Fig. 7A).** For additional control, terminally differentiated ECs (EC d6) were treated with the three substances and ZO1 assembly was analyzed 2 days later. The results of this experiment showed clearly visible gaps between ZO1-containing areas both after Dox, B02 and EST treatment **(Fig. 7A),** indicating disturbed ZO1-related tight barrier functions resulting from drug treatment of differentiated EC d6. Based on the data we conclude that Dox sensitive differentiation-associated mechanisms interfere with the formation/assembly of ZO1 containing tight junctions. At the same time, the maintenance of ZO1-containing tight junctions is impaired in terminally differentiated EC following treatment with Dox and DDR-modifying compounds. Based on this data, we conclude that substances interfering with genomic stability can (i) damage the further differentiation process of endothelial progenitor cells, thereby leading to a dysfunctional differentiated progeny and, furthermore, (ii) can impede the functionality of terminally differentiated EC. The observed drug-induced effects are independent of ZO1 (*Tjp1*) mRNA expression **(Fig. 7C)**. Analyzing the mRNA expression of the ZO1 interacting factors Claudin 5 (*Cldn5*) and Ocludin (*OcIn*), B02 treatment reduced *Cldn5* mRNA expression in EC d4, but did not affect its mRNA expression in EC d6. Neither Dox nor EST exposure of both cell types resulted in any significant alterations in the mRNA expression of *Cldn5* and *OcIn* **(Fig. 7C)**. In conclusion, we hypothesize that Dox treatment of EC d4 interferes with the development and assembly of components of the tight junction complex leading to barrier dysfunction of terminally differentiated EC d6 by post-transcriptional mechanisms. Noteworthy, drug treatment can also result in a loosening of tight junctions, leading to intercellular gaps, which likely increases the permeability of the endothelium. For additional control, the formation of cell-cell junctions involving VE-cadherin was analyzed. As monitored by immunohistochemistry-based method, VE-cadherin protein expression and localization within the cytoplasm were impaired in EC d6 differentiated from Dox-treated EC d4 **(Fig. 7B)**. VE-cadherin containing cell-cell junctions were expressed only fragmentated with VE-cadherin being distributed throughout the cytoplasm. B02 and EST treatment did not trigger defects in VE-cadherin-related cell-cell junction formation. Taken together, the data indicate that Dox-induced injury in EC d4 disturbs the formation of an endothelial barrier by terminally differentiated endothelial cells via interfering with multiple cell-cell adhesion related mechanisms.

**Figure 7:**
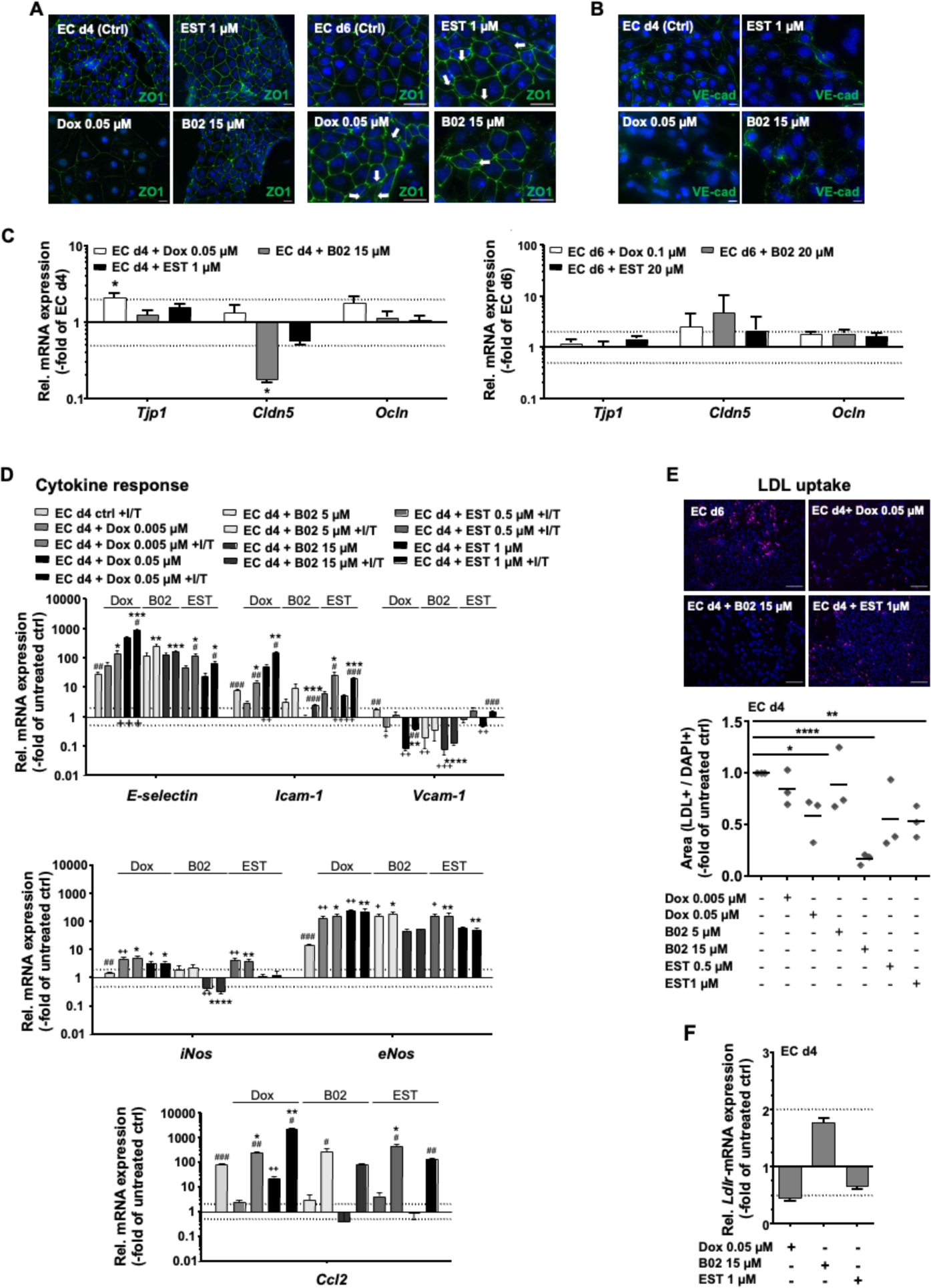
Treatment of EC d4 continuing differentiation leads to functional impairments of differentiated EC d6. **A, B:** Representative immunocytochemical staining of intracellular tight junction protein ZO1 (A) and (B) adherens junction protein VE-cadherin (VE-cad) following treatment of EC d4 and EC d6 with Dox and pharmacological inhibitors (see Fig. 1A) with the indicated concentrations. White arrows indicate gaps within the cell cluster. Scale bar: 10 µm. **C:** mRNA expression analysis of tight junction complex genes (*Tjp1, Cldn5, Ocln*) by RT-qPCR in EC d4 and EC d6 after drug exposure. EC d4 and EC d6 were treated with Dox, B02 and EST as described before (Fig 4A treatment scheme). Relative gene expression of the corresponding untreated control was set to 1.0. Only changes in mRNA-expression ≥ 2.0 and ≤ 0.5 were considered as biologically relevant (dashed lines) One-way ANOVA: p ≤ 0.05* as compared to untreated control. **D:** EC d4 were treated during ongoing differentiation for 48 h with 2 doses of Dox, EST or B02 before analyses were performed in EC d6. EC d6 were either left untreated or were exposed to pro-inflammatory cytokines (IL-1β / TNFα (+ I/T) (10 ng / ml, 1 h)) followed by mRNA expression analysis of prototypical EC-related genes *E-Selectin, Vcam-1, Icam-1, Ccl2, eNos and iNos* via RT-qPCR. Relative mRNA expression of the respective genes in untreated EC d4 control was set to 1.0. Data represent the mean ± SEM of three independent experiments each performed in biological duplicates. The dashed lines indicate changes in mRNA levels of ≥ 2.0 and ≤ 0.5, which are considered as biologically relevant. Student‘s t–test: p ≤ 0.05*, p ≤ 0.01**, p ≤ 0.001***, p ≤ 0.0001**** as compared to EC d4 ctrl +I/T (*), treated vs. untreated (^+^), + I/T vs. – I/T (^#^) **E:** The outcome of drug treatment of EC d4 on the uptake of acetylated LDL (magenta) in terminally differentiated EC d6 was analyzed by immunocytochemistry. For quantification, the LDL positive area was calculated in relation to the DAPI counterstained area (blue). Data show the mean of n=3 independent experiments normalized to the untreated control which was set to 1.0. One-way ANOVA: p ≤ 0.05*, p ≤ 0.01**, p ≤ 0.0001**** as compared to untreated control. Scale bar: 50 µm **F:** RT-qPCR gene expression analysis of *Ldlr mRNA expression* following Dox, B02 and EST treatment of EC d4 for 48 h. Relative *Ldlr*-mRNA expression in untreated control was set to 1.0. Data show the mean ± SEM from two independent experiments each performed in biological duplicates. Dashed lines indicate changes in mRNA expression of ≥ 2.0 and ≤ 0.5 are considered as being biologically relevant.

Next, the response of EC d6 that have been differentiated from untreated or drug-treated EC d4 cells to pro-inflammatory cytokines was investigated. To this end, the mRNA expression of a subset of inflammation-associated marker genes was analyzed. EC d6 derived from non-treated EC d4-derived EC d6 were found to respond to treatment with the pro-inflammatory cytokines IL-1β and TNFα with the upregulation of the mRNA expression of adhesion molecules (*E-selectin, Icam-1*), endothelial NO-synthase (*eNos*) and the chemokine CCl2 (*Ccl2*) **(Fig. 7D)**, reflecting a healthy endothelial-like phenotype as anticipated. Dox treatment of EC d4 resulted in a profound increase in the basal mRNA expression of the adhesion factors *E-selectin* and *Icam1* in the absence of IL1β/TNFα of up to 400-fold of the untreated control and a decreased *Vcam1* gene expression in EC d6 **(Fig. 7D)**. Exposure of EC d6 derived from Dox treated EC d4 cells to IL-1β and TNFα further significantly increased the mRNA expression of the aforementioned endothelial genes **(Fig. 7D)**. Additionally, treatment with pharmacological inhibitors causes similar changes in the basal mRNA expression of the examined adhesion molecules **(Fig. 7D)**. By contrast, basal mRNA expression of the chemokine *Ccl2* was increased by treatment with Dox, but remained unaffected by B02 or EST. Notably yet, IL-1β and TNFα treatment resulted in a further substantial upregulation of *Ccl2* gene expression in EC d6 that were differentiated from drug-treated EC d4 **(Fig. 7D)**. After Dox and EST exposure of EC d4, the resulting EC d6 showed increased basal mRNA expression of *Nos* (*iNos, eNos*) as compared to the untreated condition. By contrast, treatment with B02 caused downregulation of basal *iNos* mRNA expression, while *eNos*-mRNA level was upregulated. Additional cytokine stimulation of differentiated EC d6 obtained from drug exposed EC d4 did not affect the mRNA expression levels of *Nos* in any of the cases. Overall, our data show that treatment of EC d4 with all three substances promotes the development of a distinct (agent specific) pro-inflammatory phenotype in differentiated EC d6, which is further aggravated upon cytokine stimulation as reflected by a potentiated mRNA expression of cell adhesion factors and *Ccl2*.

Moreover, receptor-mediated uptake of Low density lipoprotein (LDL) was comparatively investigated in EC d6 generated from untreated and drug-treated EC d4. Following Dox and inhibitor treatment of EC d4, a significantly reduced LDL uptake was observed in the differentiated EC d6 progeny, with B02-exposure of EC d4 revealing the strongest functional defect in the differentiated progeny **(Fig. 7E).** This functional impairment is independent of drug-induced changes in LDL receptor (*Ldlr)* mRNA expression of EC d4 cells **(Fig. 7F)**. Thus, we hypothesize that both Dox-induced damage and inhibition of DNA repair interferes with posttranscriptional mechanisms affecting the formation, internalization and/or signaling of LDL-receptor, eventually resulting in a defective uptake of LDL.

## Conclusion

Our data support the hypothesis that Dox-induced DNA damage in drug sensitive endothelial progenitor cells substantially interferes with a healthy development of specific cellular functions of the thereof derived terminally differentiated endothelial-like cells **(Fig. 8, Grapical abstract).** Of note, of mechanisms of DNA repair and DDR by the RAD51 inhibitor B02 and the HDACi EST also caused agent-specific impairments in the differentiation dependent development of endothelial functions in the surviving progeny. This finding highlights the relevance of an adequate basal DNA repair and DDR capacity, which is of relevance for adequate handling of endogenously formed DNA damage for healthy endothelial development. To summarize, based on our data, we speculate that the chronic cardiotoxicity observed in patients undergoing anthracycline-based chemotherapy may be partly driven by error-prone regenerative processes of drug damaged surviving endothelial progenitor cell population, eventually resulting in compromised functionality of the differentiated progeny and imbalanced endothelial homeostasis.

**Figure 8:**
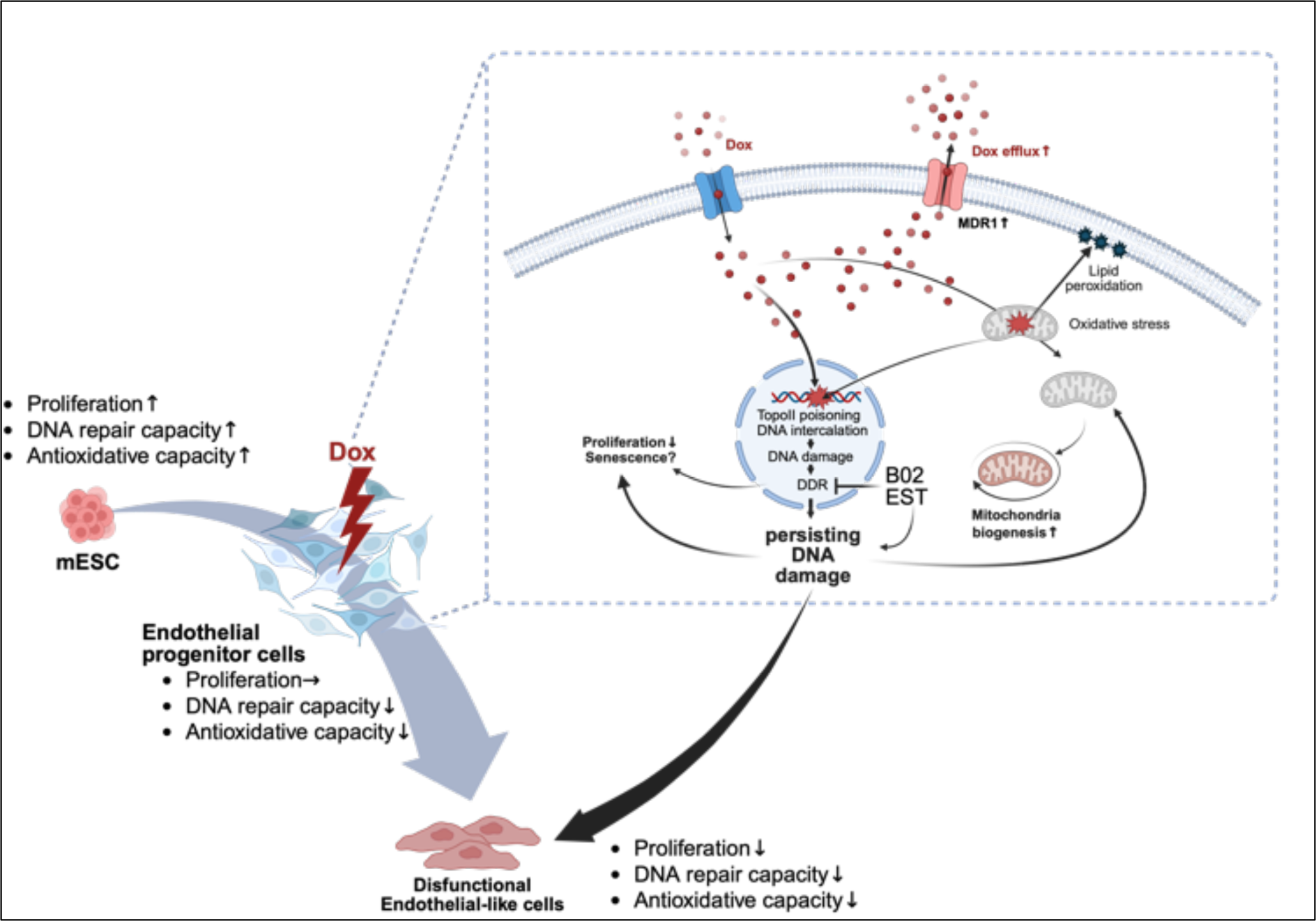
Graphical model: Dox induced DNA damage in sensitive endothelial progenitor cells resulting in compromised functionality of the differentiated EC progeny. Based on our data, we hypothesized that there is a specific Dox-vulnerable window concerning the progenitor state during endothelial differentiation that interferes with the development of physiological cellular function of the differentiated EC. This is likely due to differentiation dependent changes in DNA repair and DDR capacity, cellular detoxification and metabolism. The loss of central DNA repair factors induced by B02 and EST treatment also leading to defects of the derived EC in an agent-specific manner, highlighting the role of a robust basal DNA repair capacity in healthy endothelial development and homeostasis (created with BioRender.com).

## Acknowledgements

We would like to thank Lena Abbey for excellent technical support and Julia Mann for establishing the DNA fiber spreading assay.

## Funding Source

This work was supported by the Deutsche Forschungsgemeinschaft (DFG, German Research Foundation – 417677437/GRK2578) (subprojects: G. Fritz and A. S. Reichert).

## Author contributions

SF: Conceptualization, methodology, data generation, formal analysis, visualization, writing original draft

MW: Methodology, data generation, editing manuscript

AR: Funding acquisition, editing manuscript

GF: Conceptualization, funding acquisition, supervision, writing original draft

## Notes

### Competing Interest Statement

The authors have declared no competing interest.

### Summary of Updates

In the revised submission, flattened images were used for Fig. 1-7 to ensure correct text/chart placement in the optimized PDF provided on the hosting site.

