## Supplementary Table 1 - Primer Sequences for "Differential responses of murine embryonic stem cells (mESC) and their endothelial progeny to doxorubicin and pharmacological inhibitors of DNA repair and DNA damage response": Federmann et al-Supplementary Table 1.docx (from another version of this preprint: version 1, 2025-05-19)

| Target | | NCBI reference sequence | Sequence (5'-3') | |
| --- | --- | --- | --- | --- |
| *Abcc1* (Mrp1) | ATP Binding Cassette Subfamily C Member 1,  Multidrug Resistance Associated Protein 1 (MRP1) | NM_001425178 | fwd | GTGACTCTCAAGGGCTCCG |
|  |  |  | rev | GGGCACAGGCTTCCATAACT |
| *β-Actin* | Actin, beta | NM_007393 | fwd | GCATTGCTGACAGGATGCAG |
|  |  |  | rev | CCTGCTTGCTGATCCACATC |
| *Atp7a* | ATPase copper transporting alpha | NM_001109757.2 | fwd | ACCACCATCGCATTTGCCTA |
|  |  |  | rev | GGAGGTTTTGCCCTTCGCTA |
| *Atp7b* | ATPase copper transporting beta | NM_007511.3 | fwd | AAGAGGCCAGTCGGAAAATCT |
|  |  |  | rev | CCCCCTTCGTAGCCAACATT |
| *Brca1* | BRCA1 DNA repair associated | NM_009764 | fwd | TTGTGAGCGTTTGAATGA |
|  |  |  | rev | ACCTGGCTTAGTTACTGT |
| *Ccl2* | Chemokine (C-C motif) ligand 2 | NM_011333 | fwd | ATGAGATCAGAACCTACAACT |
|  |  |  | rev | TCCTACAGAAGTGCTTGAG |
| *Chk1* | Serine/Threonine-Protein Kinase Chk1 | NM_001410205 | fwd | GTGCCTATGGAGAAGCCAGAC |
|  |  |  | rev | CCGAAATACCGTTGCCAAGC |
| *Cldn5* | Transmembrane, tight junction protein claudin 5 | NM_013805.4 | fwd | AGTTAAGGCACGGGTAGCAC |
|  |  |  | rev | GTACTTCTGTGACACCGGCA |
| *Ctr2* | Cationic Amino Acid Transporter 2 | NM_001044740 | fwd | GTGCAGTGGCAGCTGTGAT |
|  |  |  | rev | AGCCAGGTTGGTACCTGAGA |
| *E-selectin* | Selectin, endothelial cell | NM_011345 | fwd | TGCGAGAAGAACGGATAGA |
|  |  |  | rev | CTGAATTGCCACCAGATGT |
| *Flk1* | Fetal liver kinase 1 | NM_001363216 | fwd | GTTCTGCGTGGAGACCCGAGC |
|  |  |  | rev | CAAAGCCAGTCCAGGTCCCGC |
| *Gapdh* | Glyceraldehyde-3-phosphate dehydrogenase | NM_008084 | fwd | GACGGCCGCATCTTCTTGTGC |
|  |  |  | rev | CACTGCAAATGGCAGCCCTGG |
| *Gpx1* | Glutathione peroxidase 1 | NM_008160 | fwd | TTGGTGATTACTGGCTGC |
|  |  |  | rev | TGATATTCAGCACTTTATTCTTAGTAG |
| *Gpx4* | Glutathione peroxidase 4 | NM_008162.4 | fwd | CTGGGAAATGCCATCAAAT |
|  |  |  | rev | GTCCTTCTCTATCACCTGG |
| *Gstm1* | Glutathione S-Transferase Mu 1 | NM_010358 | fwd | ACACAGCCTTCATTCTCC |
|  |  |  | rev | AATTCTAGGAAGCGTGAGTT |
| *Hmox1* | Heme oxygenase (decycling) 1 | NM_010442 | fwd | CCAGAGTCCCTCACAGAT |
|  |  |  | rev | CCCAAGAGAAGAGAGCCA |
| *Icam-1* | Intercellular adhesion molecule 1 | NM_010493 | fwd | TGCTCAGGTATCCATCCAT |
|  |  |  | rev | GGAAACGAATACACGGTGAT |
| *Ldlr* | Low density lipoprotein receptor | NM_001252658.1 | fwd | CCAATCGACTCACGGGTTCA |
|  |  |  | rev | ACAGTGTCGACTTCTCTAGGC |
| *Mdr1 (Abcb1a)* | Multi drug resistance protein 1 | NM_011076.3 | fwd | TGGAACTTGAAGAGGACC |
|  |  |  | rev | GCATAACGAAACATTGTAAGC |
| *Nanog* | Nanog homeobox | NM_001289828 | fwd | AGAGGAAGGGCGAGGAGA |
|  |  |  | rev | TTCTTGCTTACAAGGGTCTGC |
| *eNos* | nitric oxide synthase 3, endothelial cell | NM_008713 | fwd | GCATGGGCAACTTGAAGA |
|  |  |  | rev | AGGGTGTCGTAGGTGATG |
| *iNos* | Nitric oxide synthase 2, inducible | NM_010927 | fwd | ACGAGACGGATAGGCAGAGA |
|  |  |  | rev | GCACATGCAAGGAAGGGAAC |
| *Nqo1* | NAD(P)H quinone dehydrogenase 1 | NM_008706.5 | fwd | GGCCGATTCAGAGTGGCAT |
|  |  |  | rev | CCAGACGGTTTCCAGACGTT |
| *Nrf2* | Nuclear factor erythroid 2-related factor 2 | NM_001399226.1 | fwd | CCCAGCAGGACATGGATTTGA |
|  |  |  | rev | AGCTCATAGTCCTTCTGTCGC |
| *Ocln* | Tight junction protein occludin | NM_001360536.1 | fwd | GTGAATGGGTCACCGAGGG |
|  |  |  | rev | AAGATAAGCGAACCTGCCGAG |
| *Oct2* | organic cation transporter-2 | NM_001355767.1 | fwd | TCGTCACTGAGTTTAACCT |
|  |  |  | rev | AGAGATAGCATTGATGAGGAT |
| *Oct4* | octamer-binding transcription factor 4 | NM_001252452 | fwd | CTGCCCCCAGGTCCCCACTT |
|  |  |  | rev | AGCATCCCCAGGGAGGGCTG |
| *Pecam1* | Platelet endothelialcell adhesion molecule 1 | NM_001032378 | fwd | GCTCTGGGAACGAGAGCCACAG |
|  |  |  | rev | CTGCTTTCGGTGGGGACAGGC |
| *Pgc1α* | Peroxisome proliferative activated receptor, gamma, coactivator 1 alpha | NM_008904 | fwd | CCGAGAATTCATGGAGCAAT |
|  |  |  | rev | TTTCTGTGGGTTTGGTGTGA |
| *Rad51* | RAD51 homolog (S. cerevisiae) | NM_011234 | fwd | CAGCGATGTCCTAGATAATGTAG |
|  |  |  | rev | TTACCACTGCGACACCAA |
| *Sod1* | Superoxide dismutase 1, soluble | NM_011434 | fwd | ACCAGTTGTGTTGTCAGG |
|  |  |  | rev | TTTCTTAGAGTGAGGATTAAAATGAG |
| *Sod2* | Superoxide dismutase 2, mitochondrial | NM_013671.3 | fwd | GAATGTAATCAACTGGGAGAAT |
|  |  |  | rev | CATAGAATTATCAGGTATGTGGAA |
| *Tie1* | Tyrosine kinase with immunoglobulin-like and | NM_011587 | fwd | CAATGGGCCGTCTCCCTGTGC |
|  |  |  | rev | GCGTGCCTCCAAGGCTCACT |
| *Vcam-1* | Vascular cell adhesion molecule 1 | NM_011693 | fwd | ATATACTTGGAAGTGTCGTGTT |
|  |  |  | rev | GACCATCTTCACAGGCATT |
| *VE-cad* | Vascular endothelial cadherin | NM_009868 | fwd | AAGGCTCAGCGCAGCATCGG |
|  |  |  | rev | TCGTTGGCCTCCACAGTCAGGT |
| *Tjp1* | Tight junction protein ZO-1 | NM_001163574 | fwd | CCCGGACTTTTGTCCCACTT |
|  |  |  | rev | CCATTGCTGTGCTCTTAGCG |
